# The shape of density dependence points to regulation of population growth beyond classical resource limitation

**DOI:** 10.64898/2025.12.30.697033

**Authors:** Onofrio Mazzarisi, Rossana Droghetti, Samuel Barton, Hebe Carmichael, Luca Ciandrini, Marco Cosentino Lagomarsino, Martina Dal Bello, Lorenzo Fant, Giulia Ghedini, Jacopo Grilli, Gabriel Yvon-Durocher, Daniel C. Reuman

## Abstract

Growth slows as population density increases, influencing species coexistence, conservation, and resource management. This slowdown is widely attributed to declining per-capita access to energy and nutrients, and standard models of resource-limited growth predict superlinear declines of growth rate with density. Contrary to this expectation, we observed sublinear declines for 90% of 3,995 bacterial and eukaryotic growth curves. High-resolution experiments with *Escherichia coli* revealed a two-phase density dependence: a dominant sublinear regime that transitions to superlinear decline only very near population saturation. The sublinear pattern was invariant across initial concentrations of the carbon source, ruling out its depletion as the cause of early slowing. These results challenge the resource-limitation paradigm and suggest that population growth is widely inhibited by previously un-appreciated processes before the main resource converted into biomass is depleted, with implications for biodiversity theory, conservation, and resource management.

## Introduction

Exponential growth cannot continue long in a finite world. This fundamental constraint manifests as the decline of per-capita growth rates with increasing population density. Such density-dependent regulation plays a central role in shaping the dynamics, stability, and resilience of ecosystems across scales [1–5]. Yet the existence of density dependence is only part of the picture. The shape of the growth rate–density relationship—whether growth declines gradually from the outset or remains nearly unaffected until densities approach a ceiling—reflects the nature of the regulatory mechanisms at work and carries consequences that extend well beyond single-population dynamics.

In ecological communities, whether species stably coexist depends largely on how each species’ growth is regulated [5–7]. Coexistence is promoted when different species are limited by different processes—that is, when regulatory mechanisms are at least partly species-specific—because such differences create distinct ecological niches [8–10]. Conversely, when all species are governed by the same process, the scope for niche differentiation is limited, and competitive exclusion becomes more likely [11]. The shape of growth rate density dependence, measured at the single-species level, can therefore reveal what regulatory processes are at play, and in particular whether they extend beyond those most commonly assumed. Moreover, the specific form of density-dependent self-regulation can itself influence the stability of diverse communities [12–15].

The shape of density dependence also has direct applied consequences. Population viability analysis and extinction risk assessment depend on the assumed form of growth rate decline [16–19]: different curvatures imply different recovery rates for depleted populations and different thresholds for vulnerability to extinction. Likewise, foundational concepts in resource management, most notably maximum sustainable yield, are sensitive to the shape of density dependence [20–24]; misspecifying that shape can lead to overexploitation or misallocation of management effort.

Despite its importance, the shape of density dependence and the mechanisms that produce it remain poorly characterized. The most widely used models of population growth make specific predictions about this shape—predictions that, as we show below, are broadly inconsistent with empirical evidence.

## Results

### Superlinear density dependence is expected under classical resource limitation

According to the prevailing view, density dependence of growth arises primarily from the depletion of limiting nutrients or energy resources. A resource-limited scenario is often modeled by the Monod equation [25], for which growth rate, *λ*, is

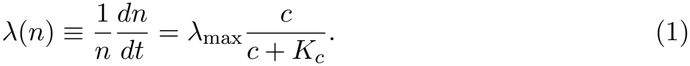

Here, *n* is population or biomass density, *λ*_max_ is the maximum growth rate attained under abundant resources, *c* is the concentration of the resource converted into biomass, and *K*_*c*_ is the half-saturation constant governing the resource level at which growth becomes limited. In a closed system, *c* decreases as *n* increases, at a rate proportional to growth

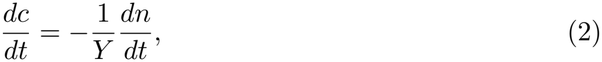

where *Y* is a yield constant. These assumptions result in a linear or superlinear dependence of *λ*(*n*) on *n* (Fig. 1 and Materials and Methods). The same conclusion holds for other more detailed growth models, such as the Droop model [26–28] (Fig. S1). One can extend consumer-resource models to include a wide range of ecological mechanisms, such as external nutrient supply, cannibalism, or co-limitation of depleted resources [29–32], and these mechanisms can, in principle, produce distinct nonlinearities in the growth-density relationships. Even so, baseline consumer-resource dynamics pre-dominantly yield linear or superlinear growth-density relationships [31].

**Figure 1:**
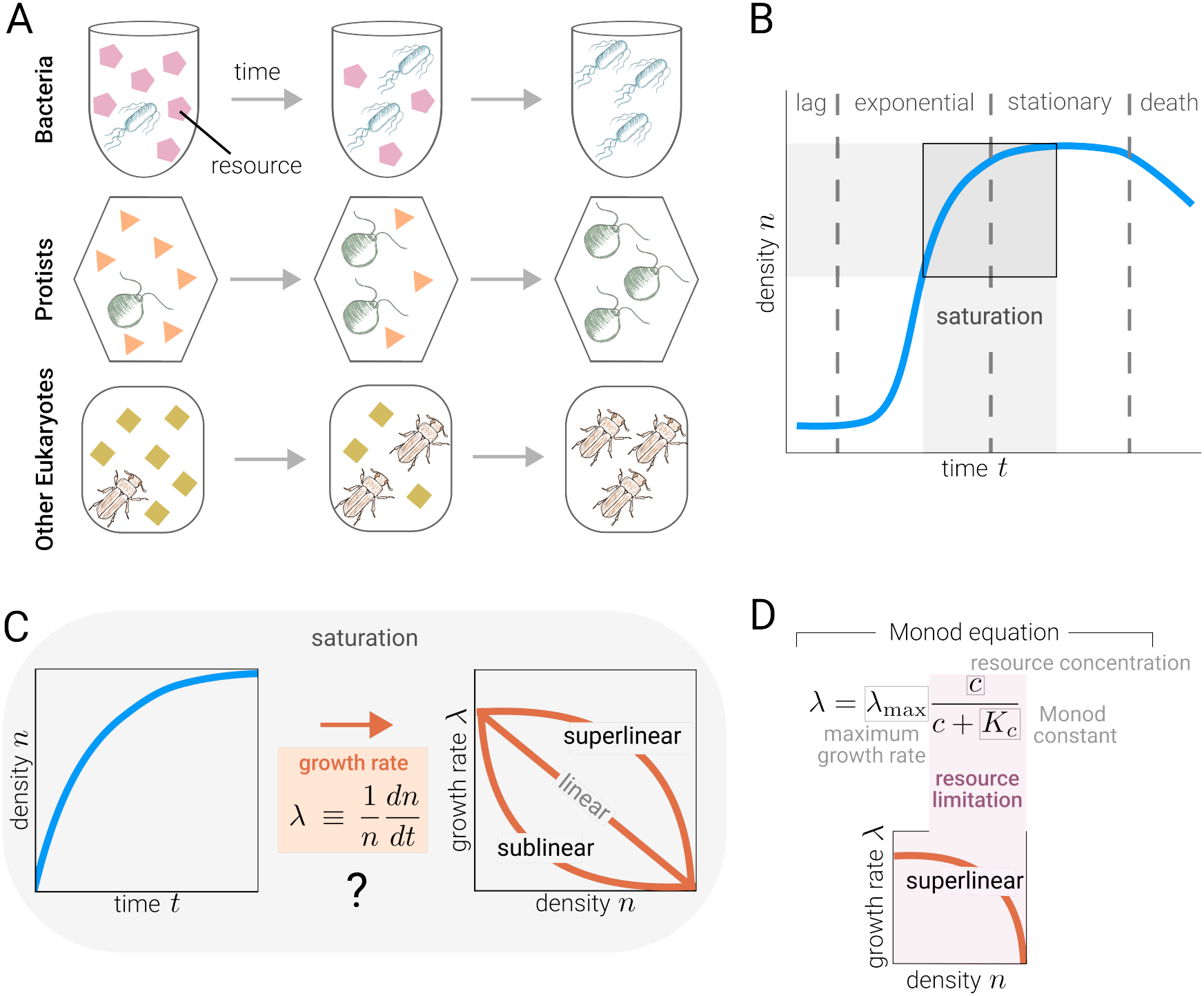
Density dependence of the growth rate is predicted to be superlinear under resource limitation. Closed lab experiments in which populations exploit a resource to grow from low to saturating densities (A) are a very common model of density dependence. Following lag and exponential phases, such systems exhibit a saturating growth phase (B) during which effective inferences can be made about the shape and mechanisms of density dependence (C). The Monod model and most other common models of population growth under scenarios of this type predict a linear or superlinear dependence of growth on density (D).

### Sublinear density dependence found across species and growth conditions

If nutrient depletion is the only phenomenon occurring as population density reaches saturation, we expect to observe widespread linear or superlinear negative density dependence. To test the generality of this prediction, we compiled a dataset of 5,871 published population growth curves covering bacterial, protist, fungal, and animal taxa, of which 3,995 passed quality filtering (Materials and Methods). We restricted our analysis to laboratory experiments in which the growth of a single isolated species was observed in a closed system initiated with a fixed amount of organisms and nutrients, and allowed to grow until the culture reached saturation (batch culture conditions, see Fig. 1 A). The only exception was a small subset of curves (dataset H, 74 of 3,995), which came from semi-continuous cultures in which part of the medium was renewed at each sampling (Table S1).

To quantify the nonlinearity of growth profiles, we used an inference tool (Materials and Methods), based on the theta-logistic model [13, 18, 33, 34]. The method estimates the theta-logistic parameters from the growth rate increments along the saturating phase of each curve via maximum likelihood. The procedure outputs an index *θ* for each growth profile, where *θ* = 1 corresponds to the linear, boundary case between sublinearity (*θ <* 1) and superlinearity (*θ >* 1) of the growth-density relationship. Our approach builds on prior efforts to quantify the curvature of density dependence [13, 17–19, 35] and improves on them in several ways (Materials and Methods), including restricting inference to the saturating phase of growth and accounting for discrete sampling, which overcome key limitations identified previously [33, 36].

Contrary to our prediction, we found that the dependence of growth rate on population density is sublinear (median *θ* = −0.36, mean *θ* = −0.34 (95% CI, −0.40 to −0.29), and 90% of growth curves had *θ <* 1) (Fig. 2). Sublinearity was consistently dominant across taxonomic groups (Fig. 2 D) and across distinct data sources and conditions (Fig. S2). Simulations confirm that the observed prevalent sublinearity in growth curves is not an artifact of the inference method (Materials and Methods and Figs. S3– S8). Per-curve profile-likelihood confidence intervals further show that this prevalence is not driven by poorly constrained estimates: nearly all curves whose interval excludes *θ* = 1 are sublinear, and sublinearity becomes more prevalent among the most precisely estimated curves (Materials and Methods and Fig. S6).

**Figure 2:**
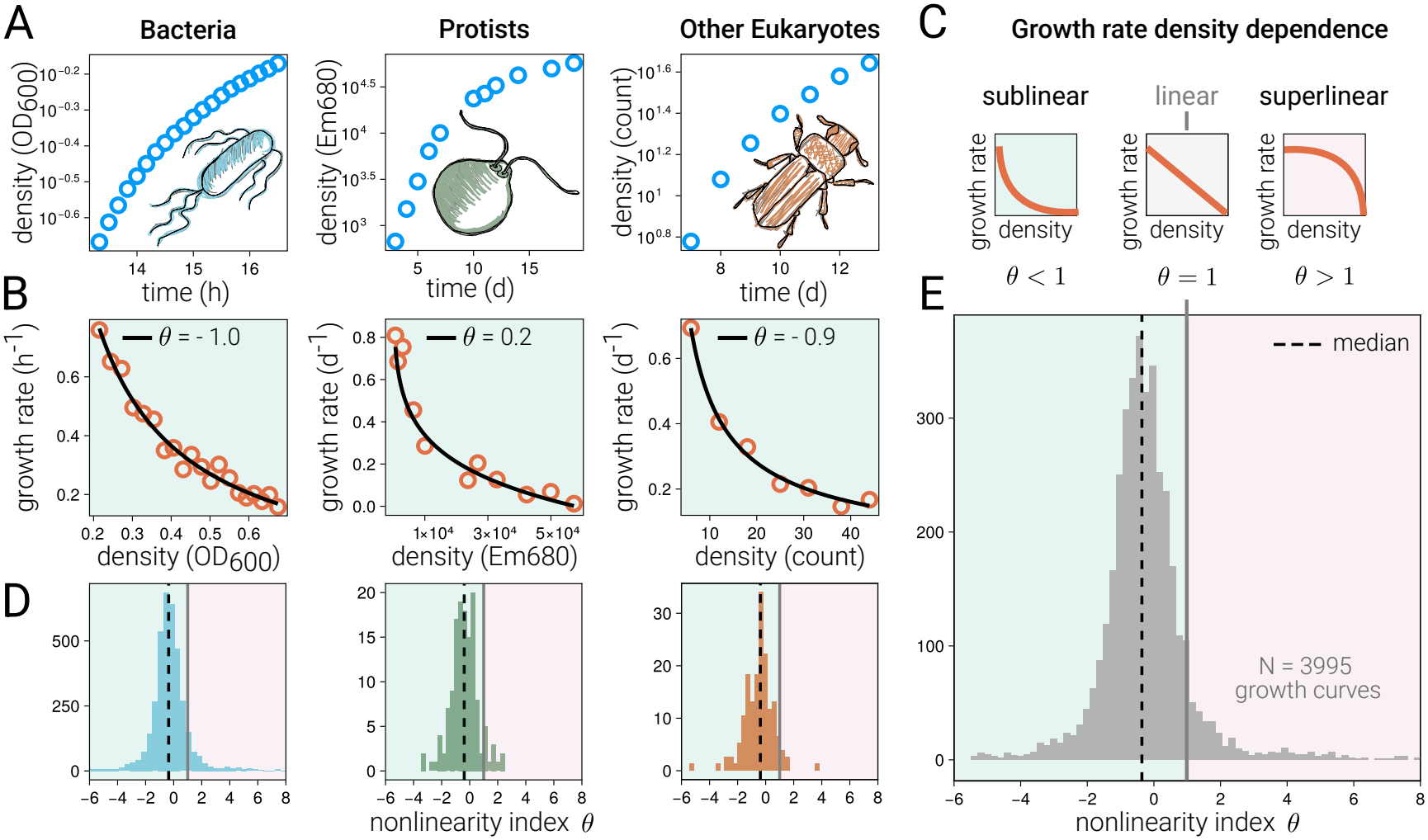
Growth rate density dependence is sublinear across species and growth conditions. Growth curves during the saturation phase (A shows examples for a bacterium (*Pseudomonas*), a protist (*Chlamydomonas*), and an animal (*Myzus*)) were converted to growth-versus-density curves (B) from which the index *θ*, interpreted as in C, was estimated (Materials and Methods). Values of *θ* were overwhelmingly less than 1, indicating sublinearity, for all three major taxonomic groups (D) and across groups (E). Vertical grey lines in D, E correspond to the boundary, linear case of *θ* = 1.

### Early sublinear growth inhibition and subsequent carbon limitation in *E. coli*

To help resolve the contrast between theory based on resource limitation and our analyses of existing growth curves, we carried out high-temporal-resolution growth experiments with *Escherichia coli*, in which population density was measured every 5 minutes across 20 technical replicates (Fig. 3 A; Materials and Methods). *E. coli* growth rates declined in a sublinear fashion at lower densities, transitioning to superlinear behavior only near carrying capacity, and thus exhibiting a biphasic structure (points in Fig. 3 B).

**Figure 3:**
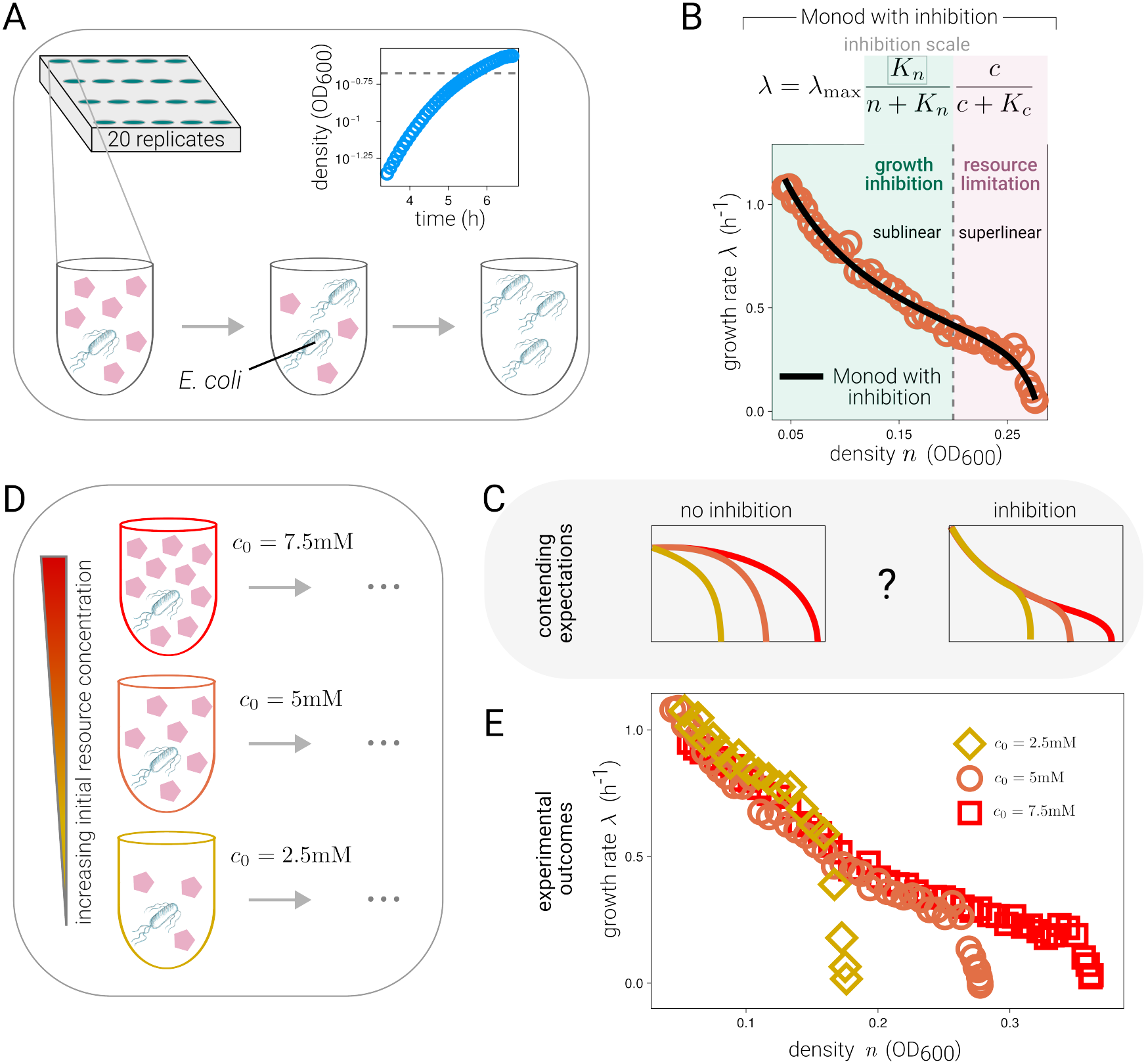
High-resolution *E. coli* experiments show biphasic structure consistent with growth inhibition independent of the carbon source concentration as a likely cause of sublinearity. (A) Schematic of the initial experimental setup and mean density across 20 replicates measured every 5 minutes. (B) Growth rate, *λ*, plotted against population density, *n*, reveals a biphasic structure characterized by early sublinear decline at low to intermediate densities, hypothesized to be associated with growth inhibition independent of the carbon source concentration, followed by superlinear decline at high densities, which matches the expected pattern for resource limitation. The black line shows the fit of the Monod model augmented with density-dependent inhibition (Materials and Methods). Shaded regions show the inhibition and resource limitation regimes. (C) Contrasting predictions: under pure resource limitation (left), the onset of growth rate decline shifts with initial nutrient concentration. Under the dual-mechanism model (right), the onset is fixed by density alone, so curves overlap during the early sublinear phase and diverge only where resource limitation takes over near the carrying capacity. (D) Schematic illustration of follow-up experiments with altered initial resource concentrations, *c*_0_ (*c*_0_ = 5mM glycerol in the initial experiment, *c*_0_ = 2.5mM and 7.5mM in the follow-ups1;8Materials and Methods). (E) Experimental growth rate density dependence profiles for different initial glycerol concentrations match the expectation of the dual mechanism model.

The biphasic structure suggests that two regulatory mechanisms co-occur: a distinct, early-acting growth inhibition that produces sublinearity at lower densities, and classic resource limitation that drives superlinear decline near carrying capacity. To formalise this hypothesis, we extended the Monod model:

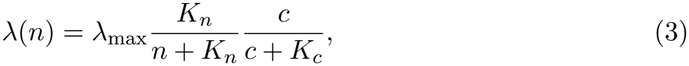

where we introduced an inhibitory term *K*_*n*_*/*(*n*+*K*_*n*_) that scales with population density. The constant *K*_*n*_ sets the population density scale at which inhibition becomes relevant, and the resource concentration *c* follows the same dynamics as in Eq. (2). For sufficiently large *K*_*n*_, our model recovers the Monod equation. The model reproduces the observed curvature shift (Fig. 3 B; Materials and Methods), which the Monod equation alone cannot generate, since it yields only linear or superlinear density dependence. The curvature of density dependence in *E. coli* appears to emerge from the interplay of two regulatory forces—inhibition and resource limitation—acting at different density scales.

To test whether the early sublinear regime is driven by inhibition independent of the resource *E. coli* grows on, we carried out additional experiments in which *E. coli* was grown at different initial resource concentrations (Fig. 3 D). In the absence of inhibition— if density dependence is dominated only by resource limitation—the density at which growth rate begins to decline depends on the initial resource concentration, so curves at different concentrations should follow distinct trajectories throughout (Fig. 3 C, left). If instead inhibition independent of the carbon source dominates the early saturating phase, curves with different initial resource concentrations should collapse onto a common sublinear profile and diverge only later, near each curve’s respective carrying capacity, where resource limitation takes over (Fig. 3 C, right). The results aligned well with the hypothesis that inhibition independent of the carbon source concentration is responsible for the early sublinear regime (Fig. 3E). Moreover, using the parameters fitted at 5 mM and only the carrying capacity measured at 2.5 mM, the model predicts the full growth-rate profile at 7.5 mM without further fitting (Fig. S9; Materials and Methods).

## Discussion

We have shown that laboratory experiments overwhelmingly display a sublinear decline of per-capita growth rate with increasing density, a pattern observed consistently across Bacteria and Eukarya, spanning diverse taxa and growth conditions, and contradicting expectations from standard resource-limitation models.

High-resolution *E. coli* experiments revealed a biphasic density dependence, with a dominant sublinear regime giving way to superlinear decline only near saturation.

Crucially, the sublinear phase was invariant across initial concentrations of the carbon source, demonstrating that the mechanisms responsible are not driven by its depletion. Because only the carbon source was varied, we cannot exclude contributions from other resources, such as oxygen or secondary nutrients, that may become limiting at high density. Only experiments of this kind, which manipulate the resource directly, can show that growth inhibition is not driven by depletion of that resource, and this cannot be demonstrated for the other datasets in our meta-analysis. However, the consistency of the sublinear signature across all major taxonomic groups (Fig. 2 D), and its contrast with the predictions of resource-limitation models, suggests that similar mechanisms, acting before the main resource converted into biomass is depleted, limit growth in other systems too. This consistency is striking because the datasets span very different resource regimes: from heterotrophic bacteria, whose growth is fueled mainly by carbon that can be taken up in many interchangeable forms, to phytoplankton, whose growth depends jointly on several non-substitutable resources such as light and nutrients.

A natural question is why the empirical observations of Fig. 2 only display the sublinear density dependence. The most likely explanation is that empirical observations are based on data with coarser temporal resolution than our high-resolution experiments, so that only the dominant of the two growth regimes is detected. The sublinear regime is likely dominant in our literature datasets (Fig. 2 D-E) because experimenters typically provide ample initial resources to ensure clear exponential phases, so that resource limitation sets in only at densities well above those at which inhibition becomes relevant. Conversely, our model predicts that superlinear density dependence should dominate when initial resources are scarce. However, scenarios in which a low-density population encounters a resource pulse are widespread in nature [37]: examples include marine phytoplankton blooms triggered when stratified surface layers are disrupted by mixing events that inject nutrients from deeper waters [38], the seasonal succession of lake plankton following spring turnover [39], and the flush of soil microbial activity that follows the rewetting of dry soils [40]. A central implication of our results is that growth inhibition acting before the resource converted into biomass is depleted, and the sublinear density dependence it produces, may be far more common in both natural and laboratory settings than previously appreciated.

While our phenomenological model captures the observed curvature shift, it does not identify the specific physiological mechanisms underlying this inhibition. Candidate mechanisms span a range from passive consequences of growth to active regulatory responses. In the first category, the accumulation of metabolic by-products, self-inhibitory compounds, or the presence of phages provides a density-dependent brake that requires no dedicated regulatory machinery [41–45]. In the second, density-sensing systems such as quorum sensing [46] could actively suppress growth at high density. In eukaryotes, conspecific chemical cues and active metabolic down-regulation can suppress growth even in nutrient-replete conditions [47, 48]. Determining which mechanisms predominate, and whether common principles emerge across taxa, is an important open question.

Our results raise the question of whether such early growth inhibition has adaptive value. At the population level, gradually slowing growth before resources are exhausted allows a larger fraction of cells to remain viable through subsequent starvation, improving recovery and resilience [49, 50]. At the community level, widespread self-limitation could increase resilience to perturbations or environmental transitions by promoting coexistence and community diversity [5, 13, 14, 51].

Finally, while we focused on batch-culture conditions, chemostats provide another setting for controlling and measuring growth [52]. In this setting, it is possible to reconstruct growth rate density dependence profiles [29, 53], and compare them to batch measurements (Materials and Methods). A recent chemostat-based study of *E. coli* reported superlinear density dependence [53]. Since chemostat and batch experiments probe the same growth rate–density relationship [29, 53] (Materials and Methods), that result is consistent with our low initial glycerol concentration case, given the density levels considered in that study (Fig. 3 E).

Overall, our results show that, in *E. coli*, growth inhibition begins before the carbon source is depleted, and suggest that inhibition acting before the main resource converted into biomass runs out is far more prevalent across species than previously recognised, leaving a characteristic sublinear signature in the density dependence of growth. Because the shape of density dependence propagates directly into predictions of species coexistence, population viability, and sustainable yield, incorporating such inhibition into ecological and management models may substantially revise our understanding of these fundamental processes.

## Materials and Methods

### Data

Our analysis combines newly generated growth data with previously published population time series spanning bacteria, protists, fungi, and animals. All datasets are publicly available in the repository associated with this manuscript.

Dataset A was generated for this study. It consists of high–time-resolution growth experiments on *Escherichia coli* grown in batch culture under controlled laboratory conditions.

Datasets B–I were compiled from published work [44, 54–59] and reanalyzed here. Table S1 summarizes their taxa, culture conditions, density measures, and numbers of growth curves, and Table S2 lists the studies composing dataset I. These datasets include laboratory and mesocosm growth experiments on freshwater and marine bacteria, unicellular eukaryotic phytoplankton and protists, fungi, and animals, and span a wide range of environmental conditions and taxonomic groups. Some of the original studies contain multi-species experiments; we included only the single-species experiments.

Datasets B-I were used for Fig. 2, while dataset A was used for Fig. 3. For Fig. 2 A, we used time series dE1 from dataset E (bacteria), dF6 from dataset F (protists), and dI19 from dataset I (other eukaryotes).

### Dataset A

#### Strains

The strain used in this study is the wild-type *E. coli* K-12 strain NCM3722 [60].

#### Growth medium

The minimal medium (M9) used in this study is composed of: (I) Salts medium containing Na_2_HPO_4_ *·*2H_2_O (44.5 g), K_2_HPO_4_ (15 g), NaCl (2.5 g), NH_4_Cl (5 g) in 200 ml of deionized water, and (II) Complementary salts medium MgSO_4_ (2 ml of 1M solution), CaCl_2_ (100 *µ*l of 1 M solution), tryptophan (1 ml of a 4 mg/ml solution) thymidine (2.5 ml of a 2 mg/ml solution) in 199.4 ml of deionized water. To make 200 ml of M9, mix 40 ml of salts medium and 40 ml of complementary salts medium with 120 ml of deionized water. The nutrient source (glycerol)was added to M9 at the prescribed concentrations for each experiment.

#### Growth protocol

For growth experiments, we applied part of the procedure described in Ref. [50, 61], except for the last growth step that here is performed in a plate reader. We recall here the main steps. Before each experiment, we took the cells from the -80^°^C stock, placed them on an LB agar plate, and grew them at 37^°^C for one day before putting the plate in the 4^°^C fridge. Growth was then carried out in three steps: seed culture, overnight culture, and experimental culture. In the first two steps, the cells were cultured at 37^°^C in a shaker at 180 rpm. The last step was carried out in BioTek Synergy-H1 multimode plate reader at 37^°^C with double orbital constant shaking. The plate reader data was collected with Gen5 - Microplate Reader and Imager Software (BioTek, version 3.03). The seed cultures are prepared using 2 mL of fresh LB medium in sterile plastic tubes and inoculated with a single colony from the stock plate. Every experiment is prepared from a single biological replicate to ensure equivalence between the wells. After 3-4 hours of growth, the seed culture is washed via centrifugation (at 10600 rpm for 4 minutes), suspended in the minimal medium, and inoculated in a new sterile 10 ml plastic tube with 5 ml of M9+ the selected nutrient source. The size of the inoculum is calculated to ensure the cells are still growing exponentially on the morning of the day after. Cells perform around ten doublings during this phase. For the third step, a sample of the overnight culture is inoculated into fresh medium of the same composition as the overnight culture and dispensed into a 48-well plate at 0.5 ml per well. The inoculum volume is chosen so that, after three doublings, the optical density reaches approximately 0.05. The plate was immediately loaded into the plate reader, and the cells were left growing for 8 to 10 hours at 37^°^C, enough time for them to consume all the nutrients and reach the stationary phase, measuring absorbance at 600 nm (OD_600_) every five minutes. For each condition, we leave one column (6 wells) without inoculum to estimate the blank. We do not include in the analysis the most external wells of the plate to avoid edge effects. Therefore, we have a total of 20 technical replicates per experiment.

### Data analysis

We use the theta-logistic model to estimate the degree of nonlinearity in the functional dependency of the growth rate on density. The model allows us to differentiate between sublinear and superlinear per-capita growth rate vs. density, respectively corresponding to *θ <* 1 and *θ >* 1. The case *θ* = 1 corresponds to linear dependence.

The theta-logistic model can be defined as

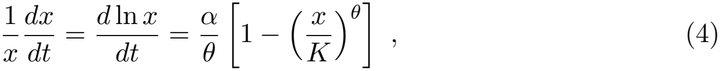

where *K* is the carrying capacity, *α* is a positive parameter that can be related to growth rate, and *θ* encodes the degree of nonlinearity in self-regulation. The presence of 1*/θ* in front of the per-capita growth rate allows for the recovery of all the nested models (including the Gompertz model for *θ* → 0) and helps reduce spurious correlation between *α* and *θ* [13, 33]. The time-dependent solution of the theta-logistic model, Eq. (4), is

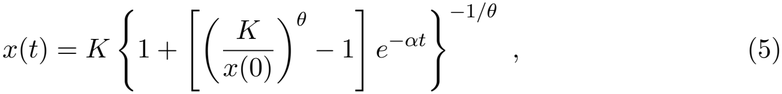

where *x*(0) is the population density at time *t* = 0.

Our data are empirical time series of population density 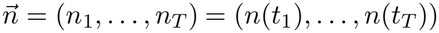, where the times of the 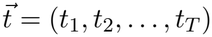 can be different for different trajectories and *T* indicates the total number of reads. We define the empirical increments as

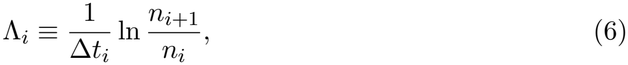

for *i* ∈ *{*1, 2, … , *T}* and where Δ*t*_*i*_ ≡ *t*_*i*+1_ − *t*_*i*_. We model the increments as

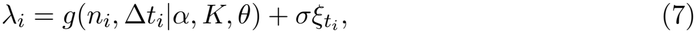

where 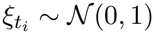 are independent and identically distributed standard normal random variables, and

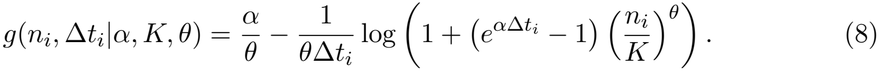

Equation (8) accounts for finite sampling intervals [36], and is obtained using the time-dependent solution (5) in the logarithmic ratio of the density at different times, divided by the time interval

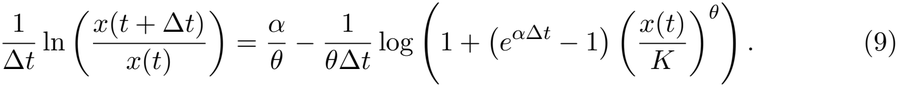

In the limit Δ*t*_*i*_ → 0, Eq. (8) reduces to

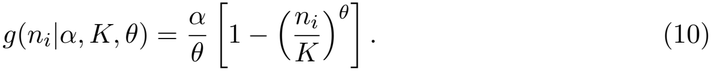

The parameters are *ρ* = (*α, K, θ, σ*), and we want to find the set maximizing the likelihood of the trajectory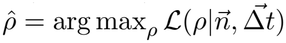, the likelihood *ℒ* being

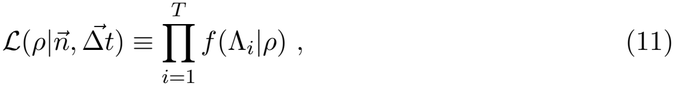

where *f*(Λ_*i*_|*ρ*) is a gaussian distribution with mean *g*(*n*_*i*_, Δ*t*_*i*_|*α, K, θ*) and standard deviation *σ*.

We are interested in the saturating part of the growth curves, and we use the following operational rationale to select the portion of the time series we consider for the fit. We selected the initial point as the one corresponding to the highest increment, representing the inflection of the density curve or the last point of the exponential growth phase. All results are qualitatively robust to instead selecting the next point or the one after that (at the cost of potentially losing informative data). We selected as the final point the last positive increment before the first negative increment, marking the onset of a declining phase or fluctuations around the stationary value. For the analysis, we only consider time series with at least four time points retained after filtering. Of the 5,871 compiled curves, 1,688 could not be fitted. Among the fitted curves, we discarded those whose estimate reached the edge of the admissible range, 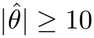 (37 at the upper and 135 at the lower bound, so that this criterion removes more sublinear than superlinear estimates), and degenerate fits with 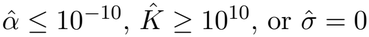 (16 curves), leaving 3,995 curves for the analysis.

To test the reliability of the method in determining the overall sublinearity or super-linearity of the growth rate density dependence, we performed a robustness analysis using synthetic data. We generated artificial population growth trajectories from the theta-logistic model with known parameters (*α, K, θ*) and additive Gaussian noise, reproducing the same statistical structure assumed for the empirical increments (Eq. (7)). Synthetic time series were sampled at finite temporal resolution, processed to compute empirical increments, and analyzed using the same maximum-likelihood inference pipeline and filtering criteria applied to the real data.

In these simulations, the nonlinearity index *θ* was sampled independently for each trajectory from a normal distribution centered on a prescribed ground-truth value, *θ* ∼ *N*(*θ*_0_, *σ*_*θ*_). This procedure mimics heterogeneity across experimental replicates and ensures that robustness is evaluated with respect to both noise and intrinsic variation in density dependence. All other parameters were held fixed within each simulation set.

We explored a broad range of parameter values, focusing in particular on the recovery of the nonlinearity index *θ* across sublinear (*θ <* 1), linear (*θ* = 1), and superlinear (*θ >* 1) regimes. As shown in Fig. S3, the inferred values of 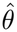 closely track the true values used to generate the synthetic trajectories, with fluctuations arising primarily from finite sampling and noise. Importantly, the inference reliably distinguishes sublinear from superlinear density dependence across the explored parameter space.

The distributions of inferred parameters are approximately unbiased and centered on the ground-truth values for *θ, α*, and *K* (Fig. S3 B–D). Additional analyses confirm that the fitted parameters do not exhibit strong mutual correlations in the synthetic data (Fig. S4), indicating that the estimation of *θ* is not confounded by trade-offs with other parameters. Together, these results reinforce the conclusion that the predominance of sublinear density dependence observed in the empirical datasets is not an artifact of noise, temporal discretization, or the inference procedure itself.

The fitted parameters do not show a strong correlation, as shown in Figs. S4 and S5, relative to simulations and real data, respectively. There is only a weak positive correlation between *θ* and *α* (Fig. S5 A) and weak negative correlation between *α* and *K* (Fig. S5 C), resulting in zero correlation between *θ* and *K* (Fig. S5 B). The same pattern is observed both in the simulated data (Fig. S4) and in the real data (Fig. S5), indicating that it is not problematic for the inference of the correct parameters, as we showed that the method is reliable with simulated data (Fig. S3).

To quantify the uncertainty of each estimate of *θ*, we computed profile-likelihood confidence intervals [62]. For each growth curve, we fixed *θ* on a grid spanning the admissible range −10 ≤ *θ* ≤ 10 and maximized the likelihood over the remaining parameters. After maximizing over *σ*, the profile likelihood depends on *θ* only through the minimal residual sum of squares RSS(*θ*) of the increments within the fitting window, and the 95% confidence interval is the set of values of *θ* not rejected by the data,

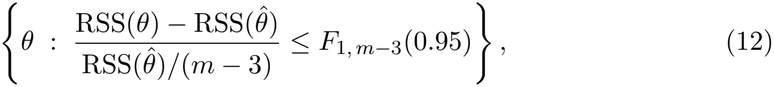

where *m* is the number of increments in the fitting window and *F*_1, *m*−3_(0.95) is the 95% quantile of the *F*-distribution with 1 and *m* − 3 degrees of freedom. We used this small-sample form instead of the asymptotic *χ*^2^ threshold because many fitting windows are short. Interval endpoints were obtained by linear interpolation between grid points, and an interval was considered unbounded when it reached the edge of the admissible range. Because consecutive increments share a density measurement, the independence assumption of Eq. (7) holds only approximately, and the intervals should be read as approximate.

The intervals for all 3,995 curves are shown in Fig. S6. About half of the curves have intervals that include *θ* = 1, especially those with short fitting windows, so individual curves often cannot discriminate between sublinear and superlinear density dependence.

However, among the 1,960 curves whose interval excludes *θ* = 1, 97.2% are sublinear (1,906 with the interval entirely below 1, against 54 entirely above). Moreover, the fraction of curves with 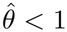 increases when the analysis is restricted to more precisely estimated curves, reaching 98.6% among curves with interval width below 2 (*n* = 1,755), while the median of 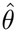 is essentially unchanged. Point estimates above 1 are instead mostly poorly constrained (58% have intervals unbounded on at least one side, against 18% for estimates below 1). The prevalence of sublinearity is therefore not driven by imprecise estimates.

### Bias in nonlinearity estimation for stationary time series

A common approach to studying the form of self-regulation involves analyzing time series of populations fluctuating around a stationary value, fitted with a theta-logistic model [18]. However, using stationary data can be problematic. We show below, analytically and through simulation, that in the case of undersampled time series, there is a bias towards *θ* → 0 (Gompertz model). The effect is more pronounced the more severe the undersampling.

Let us now denote with *n*_*t*_ the density of the population at time *t*. The increments are defined as

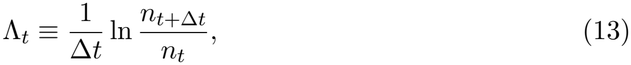

and for simplicity, we assume that samples are equally spaced in time every Δ*t* time units. We model the increments as in Eq. (7). We can always write

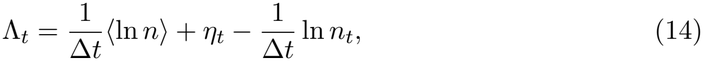

where ⟨ln *n*⟩ is the mean value of the logarithm of the density and *η*_*t*_ is an increment which can, in principle, depend on *n*_*t*_ and previous values in time. If the system is at steady-state and we sample it at a rate at which each measurement is uncorrelated with the previous one, i.e., the lag 1 autocorrelation tends to zero

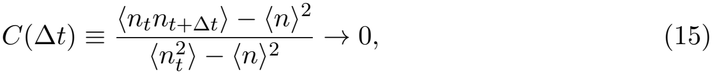

then we can approximate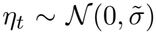, with some standard deviation 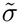. In this case, the correct generating model for the time series is

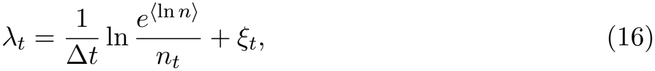

which is Gompertz model, i.e., theta-logistic with *θ* → 0, where *α* = 1*/*Δ*t, K* = *e*^⟨ln *n*⟩^ and 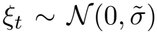. In Fig. S7, we show the effect of undersampling a stationary time series on the estimation of *θ*. In particular, we integrate the dynamics of the stochastic differential equation

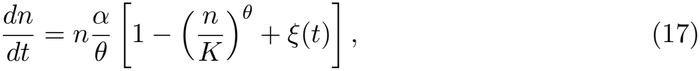

with ⟨*ξ*(*t*)⟩ = 0 and ⟨*ξ*(*t*)*ξ*(*s*)⟩ = *δ*(*t* − *s*)*σ*^2^, for a given ground truth *θ*, and we sample the solution with different frequencies.

We analyzed the Global Population Dynamics Dataset (GPDD) time series [63] in the light of the results on biases in fitting stationary time series. We find that overall, the autocorrelation of the time series in the database is not negligible (around 0.44), but we cannot exclude some level of bias in the estimation of the parameter *θ*. In Fig. S8, we compare the GPDD time series with the same database in which we shuffled each time series (i.e., for each time series, we randomly reordered the time points). The mean value of theta for the shuffled ensemble converges to *θ* → 0 as expected.

Focusing, in the present work, on time series of populations growing from low to high density in batch culture conditions, we avoid this bias as well as other problems in the use of the theta-logistic model to measure the nonlinearity of the self-regulation highlighted in Ref. [33].

### Monod equation

Monod equation [25] describes the dynamics of a population *n* growing on a resource with concentration *c*

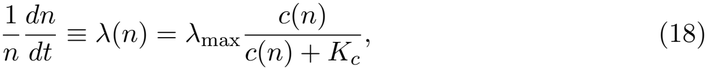

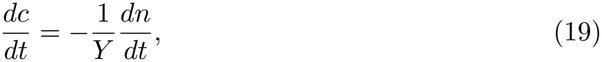

where the parameters are described in the Results section. The variation of *c* with respect to *n* reads *dc/dn* = −1*/Y* , which can be integrated to give *c*(*n*) = (*K* − *n*)*/Y* , where *K* ≡ *c*_0_*Y* + *n*_0_ is the carrying capacity, *c*_0_ the initial resource concentration and *n*_0_ the initial density. The growth rate as an explicit function of the density

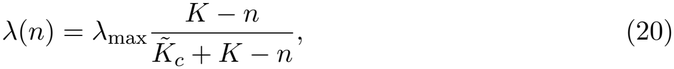

Where 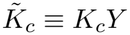, demonstrating a superlinear dependence. The dependence is approximately linear (logistic growth) as 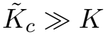.

### Monod equation with growth inhibition

The derivation of the growth rate of the Monod equation with growth inhibition with explicit density dependence is the same as for the Monod equation (20), and gives

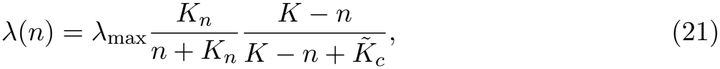

where, again,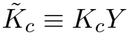. As for all growth curves in this work, the fit is restricted to the saturating phase (see Data analysis). The fitted density parameters in Fig. 3 are *K* = 0.28, *K*_*n*_ = 0.06, and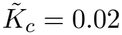, in units of optical density. The fitted *K*_*n*_ is comparable to the density at which the saturating phase begins. The fitted *λ*_max_ = 2.05 h^−1^ is the extrapolation of the inhibitory term to *n* → 0, outside the fitted range, whereas at the start of the fitted range the model gives *λ* ≈ *λ*_max_*/*2 ≈ 1 h^−1^, matching the maximal observed growth rate.

The inflection point *n*_sub→sup_, separating the sublinear phase and the superlinear phase, occurs where the second derivative with respect to *n* vanishes. Differentiating twice and setting *λ*^*′′*^(*n*) = 0 eliminates the factor *λ*_max_ and yields a cubic equation for *n*. It is convenient to shift variables via *x* = *n* − *K*, which transforms the cubic into the depressed form

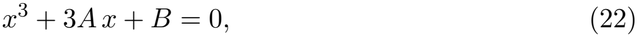

Where

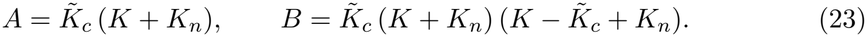

The discriminant of this cubic is

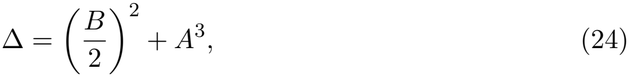

which is positive for *K, K*_*n*_, 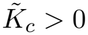, implying a single real root. The inflection point is recovered as

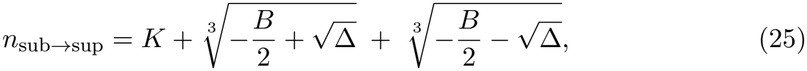

which, for the fitted parameters in Fig. 3 gives *n*_sub→sup_ = 0.2.

### Prediction for different initial resource concentrations

To test the model beyond the fitted curve, we predicted the growth-rate profile at *c*_0_ = mM using only information from the other two concentrations. Since *λ*_max_, *K*_*n*_, and 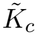 do not depend on *c*_0_, we used the values fitted at 5 mM. The carrying capacity instead depends on *c*_0_ through the yield, *K* = *c*_0_*Y* + *n*_0_. We estimated *Y* as the slope between the carrying capacity fitted at 5 mM and the final density measured at 2.5 mM, which gives *Y* ≈ 0.035 mM^−1^ (Fig. S9 A). This linear relation has a positive offset (≈ 0.10), larger than the inoculum density (≈ 0.006). Our prediction relies only on the linear dependence of *K* on *c*_0_ within the range of concentrations used, and it does not depend on the origin of this offset. Extrapolating this relation to *c*_0_ = 7.5 mM gives a carrying capacity *K* ≈ 0.36, so that no information from the 7.5 mM experiment enters the prediction, which closely matches the observed profile (Fig. S9 B).

### Comparison with chemostat framework

To compare growth rate density dependence inferred from batch cultures with results obtained in chemostat experiments, we consider a standard chemostat model in which population density *n* and resource concentration *c* evolve according to

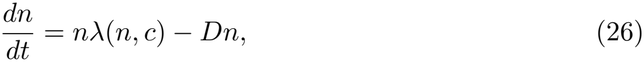

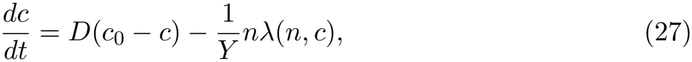

where *D* is the dilution rate, *c*_0_ is the concentration of the incoming resource, and *Y* is the biomass yield.

At steady state (*n*^*^, *c*^*^), we have

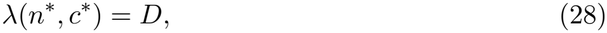

And

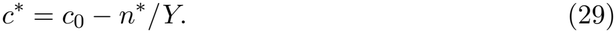

Chemostat experiments can probe the functional relationship between growth rate and population density by varying the externally imposed dilution rate *D* and measuring the corresponding equilibrium density *n*^*^ [29, 53].

For the Monod growth model,

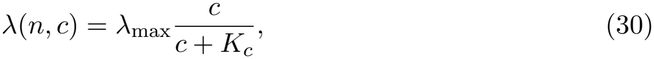

substituting the steady-state relations yields

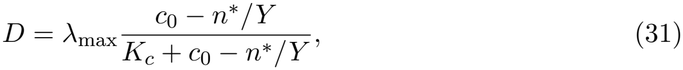

which predicts a linear or superlinear dependence of growth rate on population density, equivalent to the expectation from batch culture.

For the Monod model with inhibition,

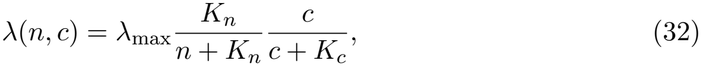

the steady-state condition instead gives

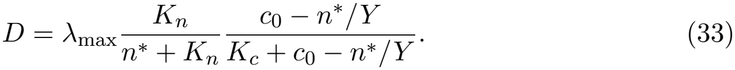

Also in this case, the same biphasic prediction obtained in batch cultures is reached.

These expressions show that chemostat and batch culture experiments probe the same underlying growth rate–density relationship, *λ*(*n*), but sample it in different ways. Batch cultures dynamically traverse a broad range of densities along a transient trajectory, while chemostats access steady-state points determined by the imposed dilution rate.

To obtain a full profile of growth rate density dependence in the chemostat setting, it is necessary to measure the steady-state density *n*^*^ for a gradient of dilution values. To explore, in a chemostat, the same phenomenology obtained in batch culture by varying the level of initial concentration (Fig. 3), it is necessary to build different profiles (each requiring a gradient of *D*) for different values of the concentration of the incoming resource *c*_0_.

## Acknowledgments

We are especially grateful to C. E. Schaum for assistance in constructing the datasets. We thank J. Aguilar, D. Amor, M. Barbier, N. Coombs, T. Cossetto, A. Goyal, G. Nicoletti, V. Karatayev, and W. Shoemaker for insightful discussions. We also thank A. Nanda for helping with the isolation of the soil bacterial strains, and J. Gore.

## Funding

O.M. acknowledges the Trieste Laboratory on Quantitative Sustainability - TLQS for funding. M.C.L. and R.D. acknowledge funding from AIRC - Associazione Italiana per la Ricerca sul Cancro AIRC IG grant no. 23258. R.D. was supported by the AIRC fellowship Italy Pre-Doc 2022 (ID 28176). L.C. acknowledges the French National Research Agency (REF: ANR-21-CE45-0009). J.G. acknowledges support by Fondo Italiano per la Scienza - FIS (CUP J53C23002290001). G.G. acknowledges support from the European Union ERC Starting Grant (META FUN, 101116029). D.C.R. was supported by U.S.A. National Science Foundation grants BIO-OCE 2023474 and DEB-PCE 2414418, as well as by the Alexander von Humboldt Foundation.

## Author contributions

O.M., G.Y.-D., and D.C.R. conceived the study. R.D. performed the experiments. O.M., R.D., S.B., H.C., M.D.B., G.G., and G.Y.-D. compiled and curated the data. O.M., R.D., L.C., M.C.L., L.F., J.G., and D.C.R. developed the theoretical framework, and O.M. analyzed the data. O.M., R.D., S.B., L.C., M.C.L., M.D.B., L.F., G.G., J.G., G.Y.-D., and D.C.R. contributed to the interpretation and conceptual framing of the results. O.M. and D.C.R. wrote the original draft, and all authors reviewed and edited the manuscript.

## Competing interests

There are no competing interests to declare.

## Data and materials availability

All data and analysis code are archived on Zenodo (DOI: https://doi.org/10.5281/zenodo.20200296) and will be made publicly available upon publication.

## Supplementary Materials

Figs. S1 to S9

Tables S1 and S2

References (64–113)

## Supplementary Materials for

**Figure S1:**
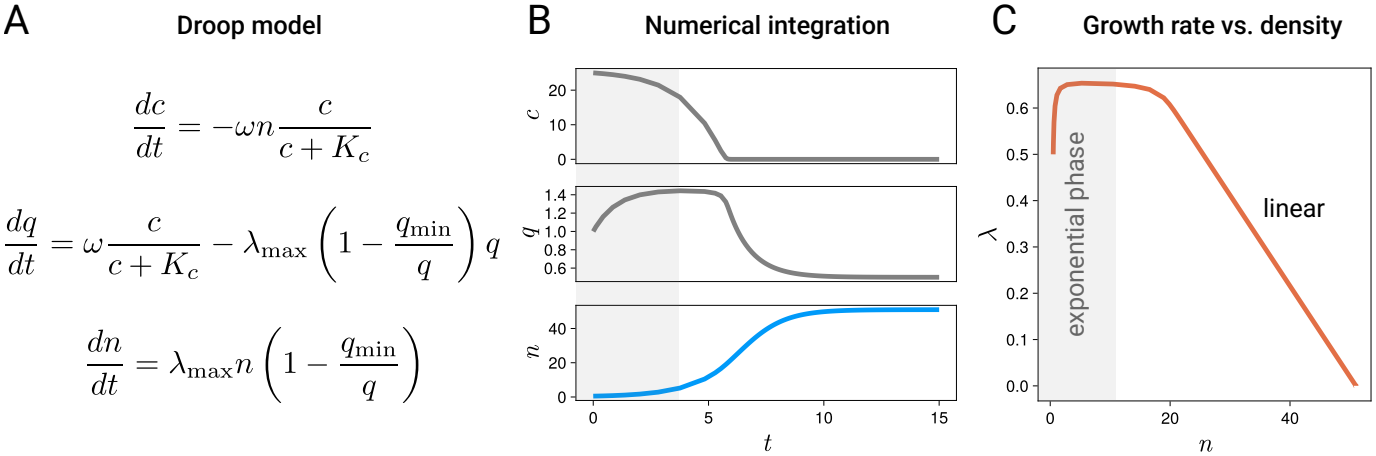
The Droop model predicts linear growth-rate density dependence under resource limitation. (A) Schematic of the Droop model, in which population growth is limited by intracellular nutrient quota. Variables: external resource concentration *c*, internal quota *q*, and population density *n*. (B) Numerical integration of the model shows depletion of external resources, relaxation of the intracellular quota toward its minimum value, and saturation of population density over time. (C) The resulting per-capita growth rate *λ* as a function of population density *n* exhibits a linear decline during the saturating phase, following an initial exponential-growth regime. This demonstrates that resource-limited growth models based on intracellular quotas do not generate sublinear density dependence. Model parameters used for the simulation are *λ*_max_ = 1, *q*_min_ = 0.5, *ω* = 1, *K*_*c*_ = 1, with initial conditions *c*_0_ = 25, *q*_0_ = 1, and *n*_0_ = 0.5.

**Figure S2:**
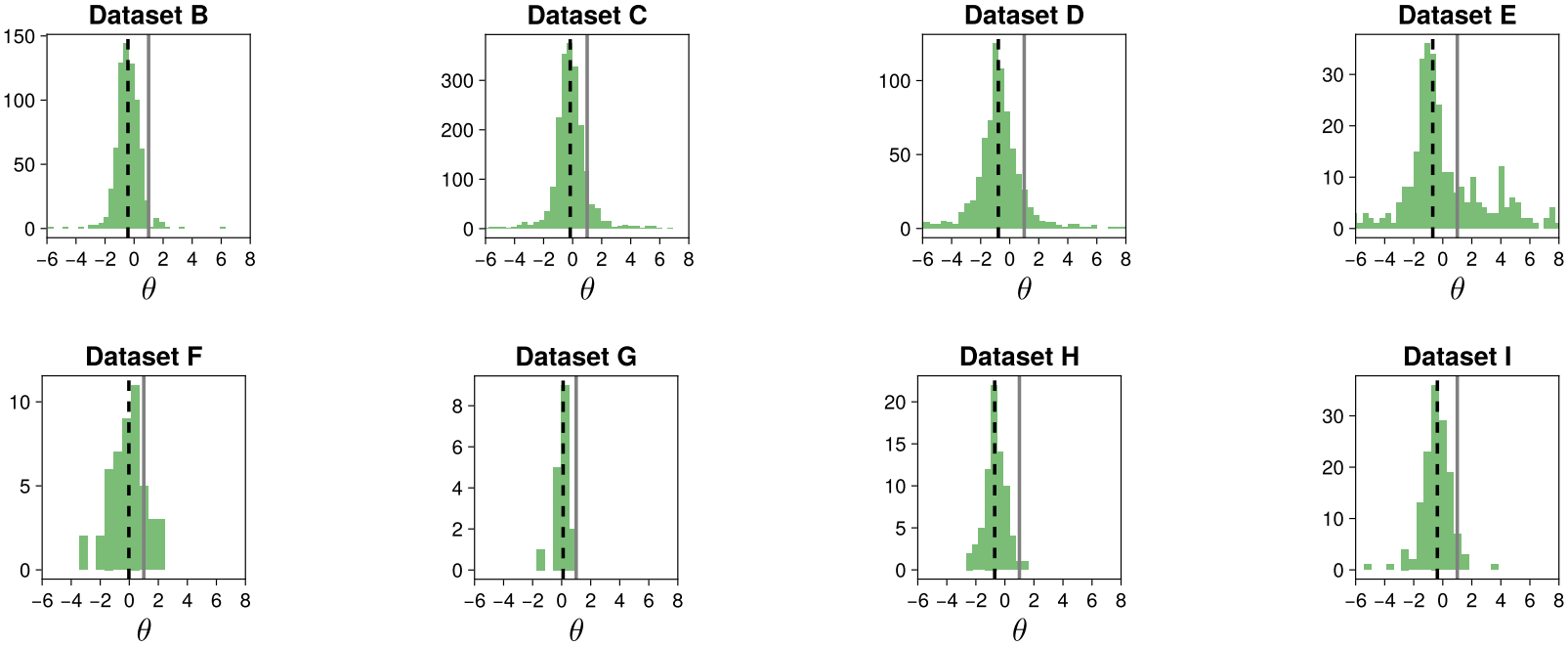
Distributions of the nonlinearity index *θ* across individual datasets. Histograms show the inferred values of the nonlinearity index *θ* for each dataset (B–I), analyzed separately using the theta-logistic inference procedure described in Materials and Methods. The vertical gray line marks the linear case (*θ* = 1), while dashed black lines indicate the median value of each distribution. Across datasets spanning different taxa and experimental conditions, inferred growth-rate density dependence is consistently sublinear.

**Figure S3:**
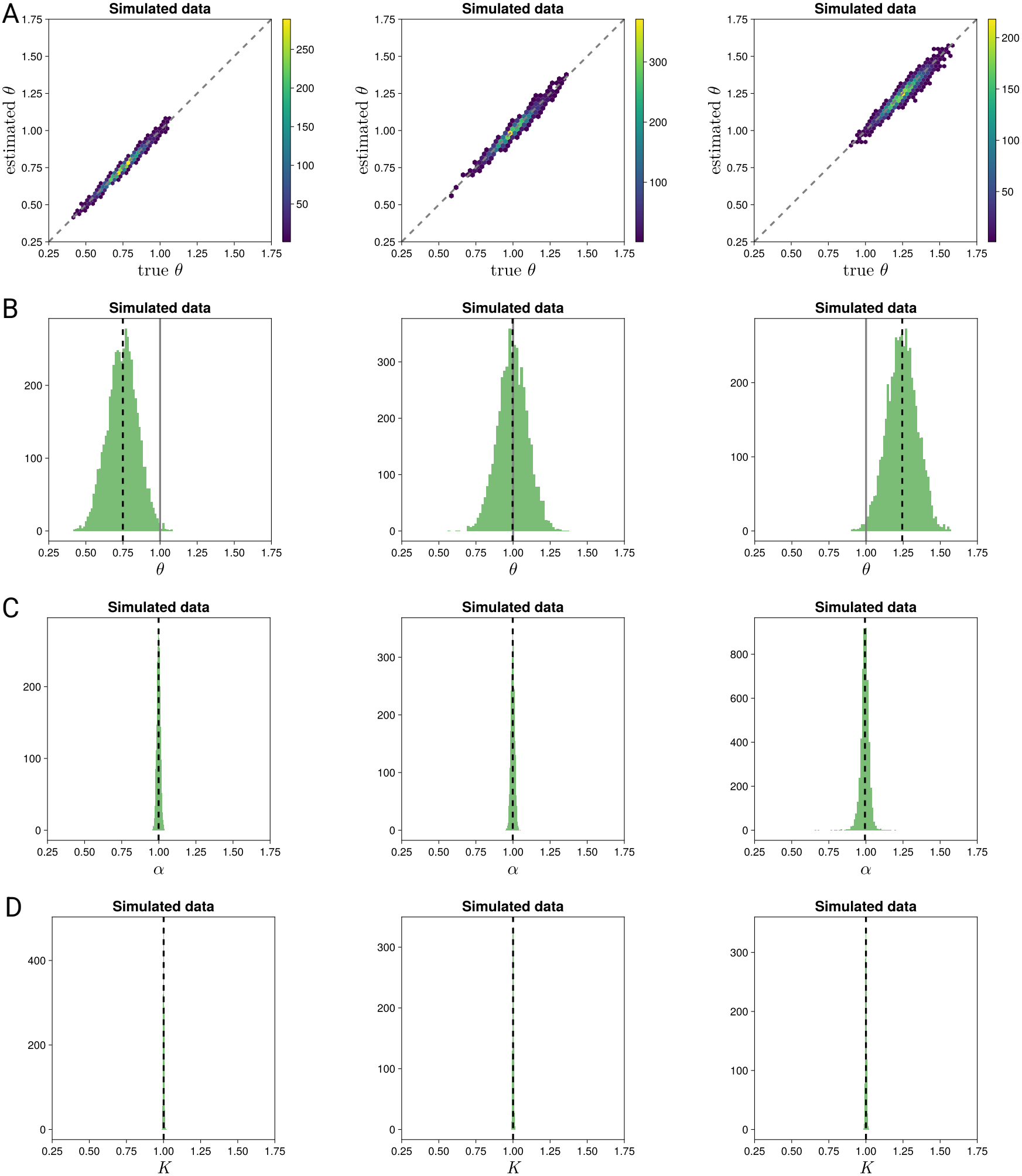
Robust recovery of theta-logistic parameters from synthetic data. 5000 synthetic trajectories were generated from the theta-logistic model with known parameters (*K* = 1, *α* = 1), with the nonlinearity parameter *θ* independently sampled for each trajectory from a normal distribution centered on prescribed values (*θ* = *{*0.75, 1, 1.25*}*) and standard deviation *σ*_*θ*_ = 0.1). For each synthetic trajectory, the initial condition is *n*(0) = 0.01, the time runs from *t* = 0 to *t* = 10, and the interval between sampled points is Δ*t* = 1. (A) Estimated versus true values of *θ* for sublinear, linear, and superlinear regimes. (B) Distribution of inferred *θ* values showing accurate recovery of the nonlinearity regime (gray line at *θ* = 1 for reference). (C) and (D) Dis-

**Figure S4:**
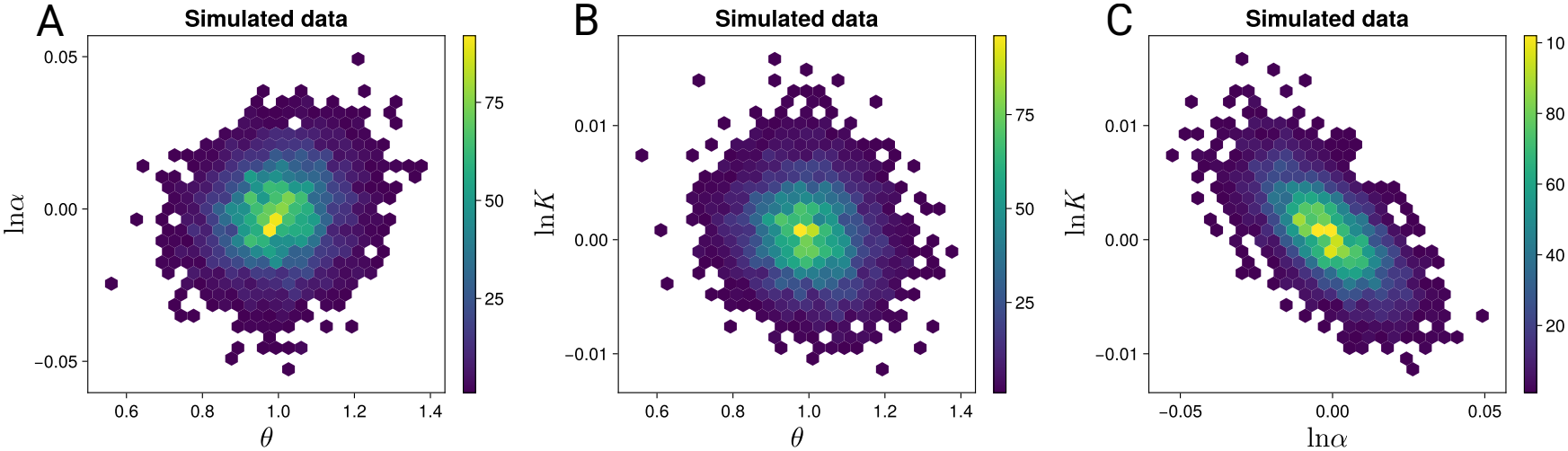
Weak correlations among fitted parameters in synthetic data. Pairwise joint distributions of inferred parameters from synthetic trajectories generated with a nonlinearity index extracted from a Normal distribution centered at *θ* = 1. (A) *θ* versus ln *α*, (B) *θ* versus ln *K*, (C) ln *α* versus ln *K*. Color intensity indicates the density of inferred parameter combinations. The absence of strong correlations indicates that parameter estimation is well-conditioned and that inference of *θ* is not driven by trade-offs with *α* or *K*. Simulation parameters as in Fig. S3.

**Figure S5:**
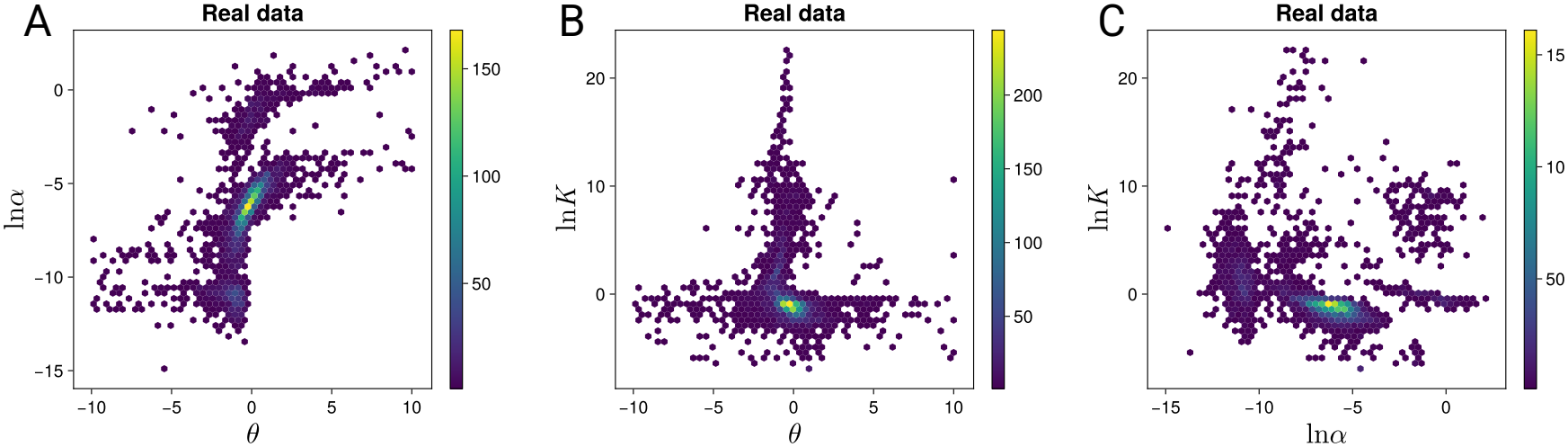
Weak correlations among fitted parameters in empirical data. Pairwise joint distributions of inferred parameters from all empirical growth trajectories. (A) *θ* versus ln *α*, (B) *θ* versus ln *K*, (C) ln *α* versus ln *K*. Color intensity indicates the density of observations. The structure closely mirrors that observed in synthetic data (Fig. S4), supporting the robustness of parameter inference and indicating that the estimated prevalence of sublinear density dependence is not driven by parameter degeneracies.

**Figure S6:**
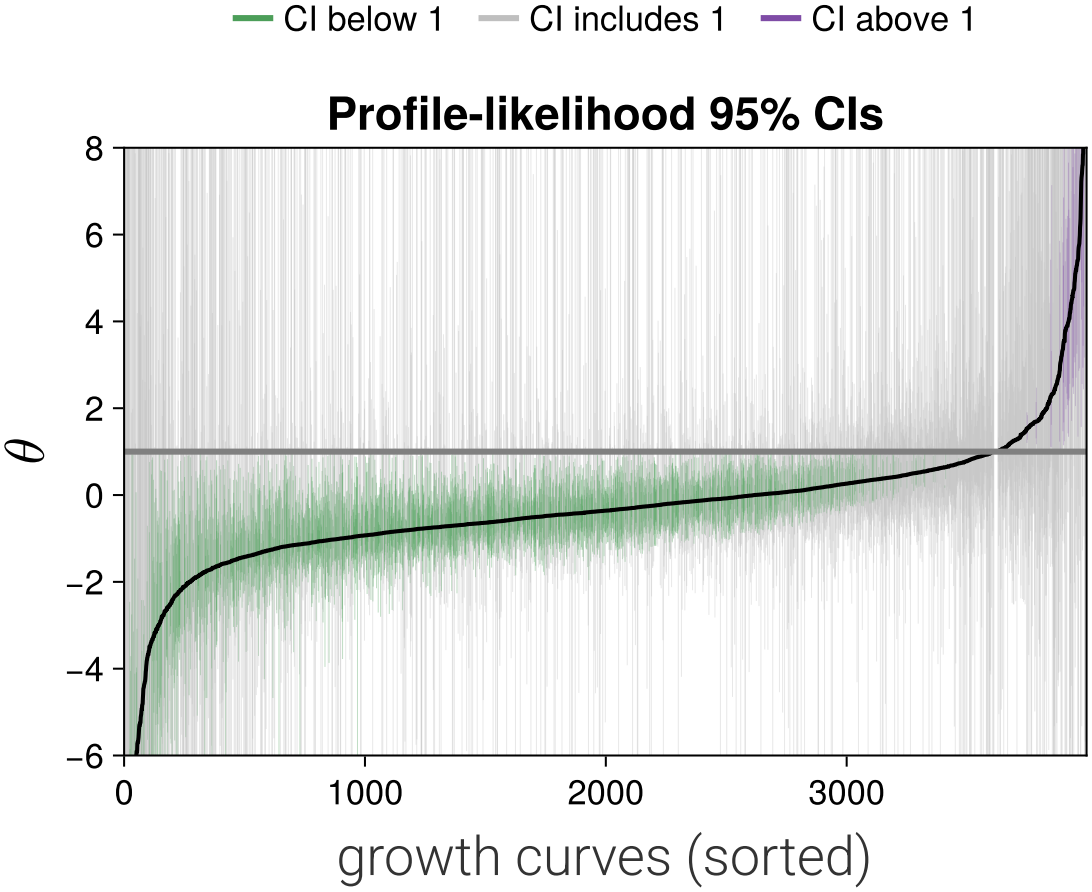
Curves whose nonlinearity is resolved are almost exclusively sublinear. Each vertical bar is the 95% profile-likelihood confidence interval of *θ* for one growth curve (*N* = 3,995); curves are sorted by the point estimate 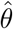 (black line). Bars are green when the interval lies entirely below *θ* = 1, purple when it lies entirely above, and grey when it includes *θ* = 1 (horizontal grey line). Intervals extending beyond the plotted range, including those unbounded within the admissible range −10 ≤ *θ* ≤ 10, are truncated at the axis limits.

**Figure S7:**
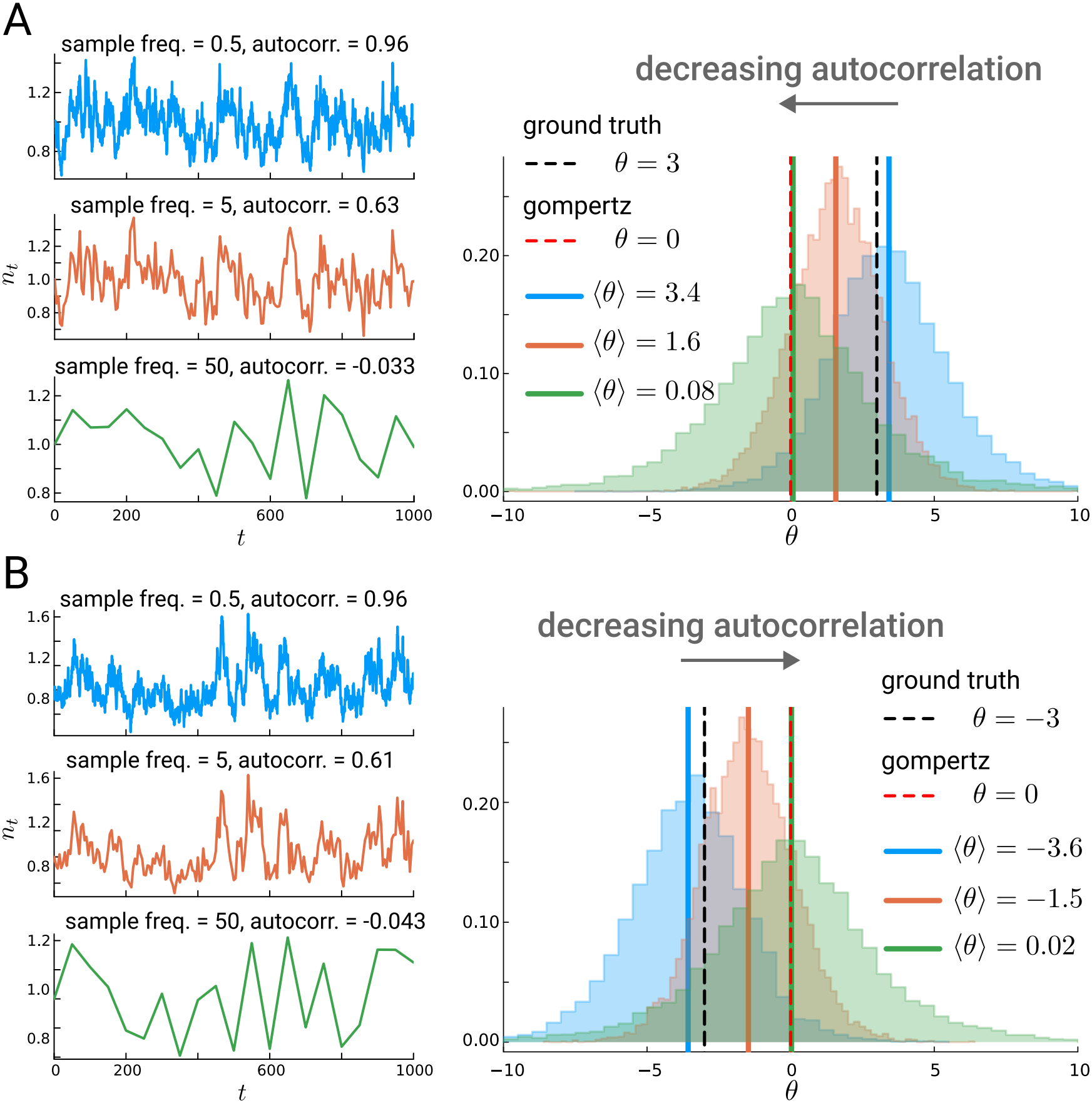
Undersampling time series at stationarity bias the estimation of *θ* towards *θ* = 0, i.e. Gompertz growth. We simulated and analyzed with our fitting procedure 10^4^ time series from *θ*-logistic model with *θ* = 3 (A) and *θ* = −3 (B). In both cases, we used *α* = 1, *K* = 1 and *σ* = 2.

**Figure S8:**
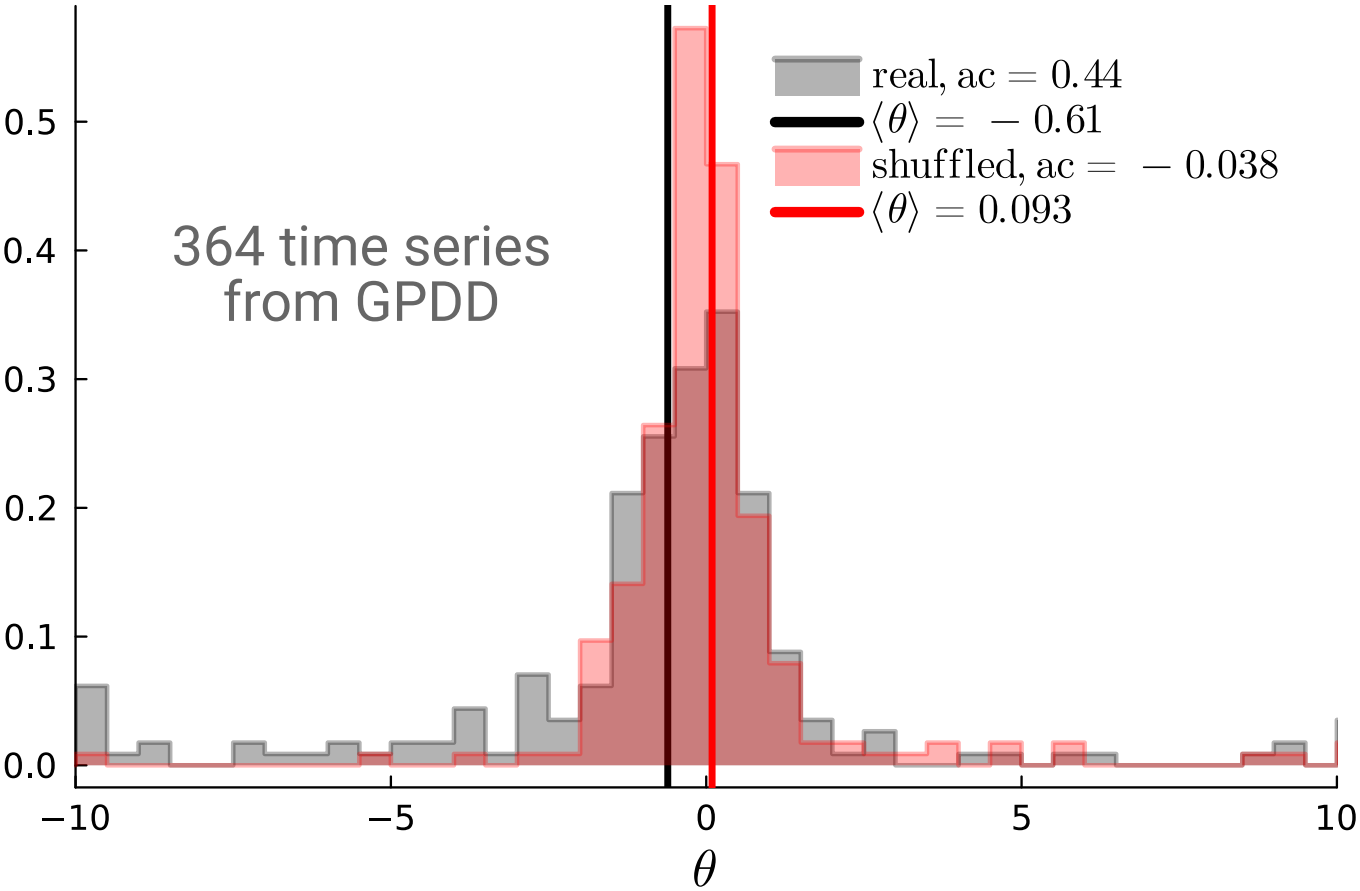
The estimated *θ* from the GPDD time series is not free from the bias towards *θ* → 0 discussed in the present work. We report here the estimated value of *θ* for 364 time series of the GPDD (gray) and for the same time series with shuffled time points (red). In the first case, we found an autocorrelation around 0.44, indicating that the bias must have an impact to some extent. The shuffled case, with autocorrelation −0.038, tends towards mean *θ* → 0 as expected.

**Figure S9:**
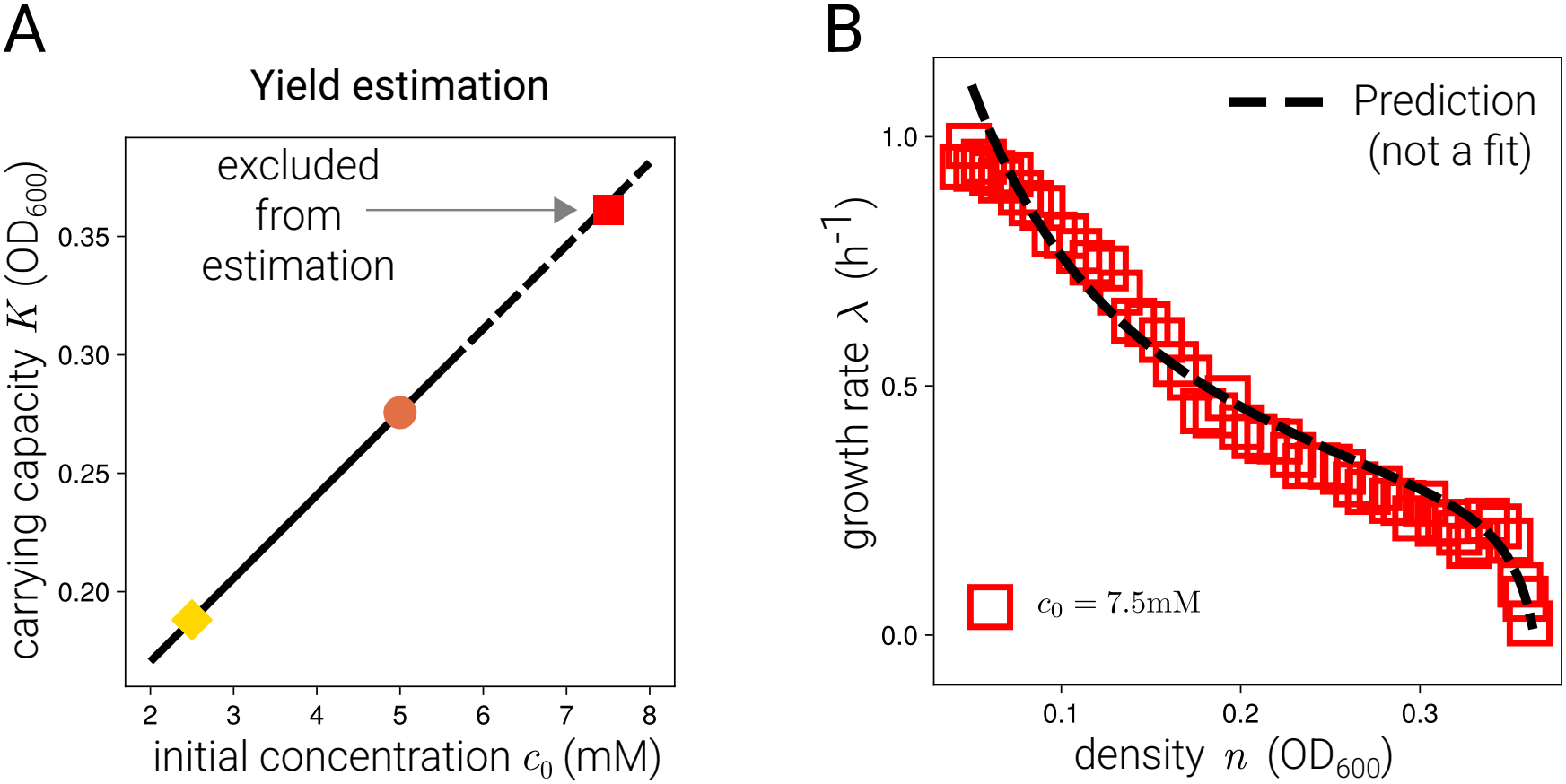
Parameter-free prediction of the growth-rate profile at *c*_0_ = 7.5 mM. (A) Carrying capacity *K* versus initial glycerol concentration *c*_0_. The line through the carrying capacities at 5 mM (circle) and 2.5 mM (diamond) defines the effective yield *Y* (slope) and is extrapolated (dashed) to 7.5 mM; the observed carrying capacity at 7.5 mM (square) is excluded from the estimation. (B) Growth rate *λ* versus density *n* at 7.5 mM (squares) and model prediction (dashed line), obtained from the parameters fitted at 5 mM (Fig. 3 B) and the carrying capacity from (A), without any fitting to the 7.5 mM data.

**Table S1:** Datasets B–I. Curves are given as retained/analyzed; retained curves are those passing the quality filters described in Materials and Methods.

|  | Taxa | Culture conditions | Density measure | Curves | Source |
|---|---|---|---|---|---|
| B | Freshwater bacteria, 36 strains (a) | Batch; temperature (15–42 °C), pH (4–11), or salinity (0–35 g/L) varied one at a time | OD<sub>600</sub> | 718/864 | [54] |
| C | Freshwater bacteria, 34 strains (a) | Batch; combinations of temperature (15–42 °C), pH (4–11), and salinity (0–35 g/L) | OD<sub>600</sub> | 1,947/2,440 | [54] |
| D | Freshwater bacteria, 12 strains (a) | Batch; combinations of temperature (15–47 °C), pH (4–11), and salinity (0–35 g/L) | OD<sub>600</sub> | 747/1,200 | [54] |
| E | Soil bacteria, 8 strains (b) | Batch; M9 minimal medium with glucose or sucrose (0.01, 0.1, or 1% w/v), with or without vitamins; 30 °C | OD<sub>600</sub> | 305/864 | [44] |
| F | Marine phytoplankton, 61 eukaryotic and 2 cyanobacterial strains (c) | Batch; 17 °C (20 °C for diatoms), 14:10 h light–dark cycle | Chlorophyll fluorescence | 48/63 | [55] |
| G | Diatom *Thalassiosira pseudonana* | Lines from constant (22, 26, 32 °C) or fluctuating temperature regimes, assayed at 15–40 °C; batch during the analyzed phase (d) | Cell counts (flow cytometry) | 17/30 | [56] |
| H | Phytoplankton: *Amphidinium*, *Tetraselmis*, *Dunaliella*, *Tisochrysis*, *Synechococcus* | Semi-continuous: 10% of the culture volume replaced with fresh medium at each sampling | Cell counts | 74/100 | [57–59] |
| I | Animals, yeast, and a cyanobacterium | Laboratory or caged populations | Counts, cell densities, biomass | 139/310 | Table S2 |
| Total |  |  |  | 3,995/5,871 |  |

(a) Isolated from Icelandic geothermal pools [64] and from experimental mesocosms in Dorset [65], identified by 16S rRNA sequencing: *Acinetobacter*, *Aeromonas*, *Arthrobacter*, *Bacillus*, *Brevibacterium*, *Burkholderia*, *Buttiauxella*, *Chromobacterium*, *Chryseobacterium*, *Curtobacterium*, *Erwinia*, *Flavobacterium*, *Frigobacterium*, *Hafnia*, *Herbaspirillum*, *Janthinobacterium*, *Kocuria*, *Microbacterium*, *Mucilaginibacter*, *Pedobacter*, *Pseudomonas*, *Serratia*, *Sphingomonas*, and *Yersinia*. (b) Isolated from serial passaging of soil communities: *Agrobacterium*, *Klebsiella*, *Paenarthrobacter*, *Pseudomonas*, *Raoultella*, *Sphingobacterium*, and *Sphingopyxis*. (c) *Amphidinium*, *Bigelowiella*, *Calcidiscus*, *Calyptrosphaera*, *Chlamydomonas*, *Chrysochromulina*, *Chrysotila*, *Coccolithus*, *Coscinodiscus*, *Diacronema*, *Dunaliella*, *Emiliania*, *Gephyrocapsa*, *Halamphora*, *Heterocapsa*, *Isochrysis*, *Karenia*, *Lepidodinium*, *Micromonas*, *Minidiscus*, *Nannochloropsis*, *Nitzschia*, *Ostreococcus*, *Pavlova*, *Phaeocystis*, *Phaeodactylum*, *Prorocentrum*, *Prymnesium*, *Rhodomonas*, *Ruttnera*, *Scrippsiella*, *Scyphosphaera*, *Skeletonema*, *Synechococcus*, *Thalassiosira*, and *Thoracosphaera*. (d) Cultures were transferred to fresh medium during exponential growth and then left to reach stationary phase. Because no medium was added after the last transfer, the saturating phase analyzed here corresponds to batch-culture conditions.

**Table S2:**
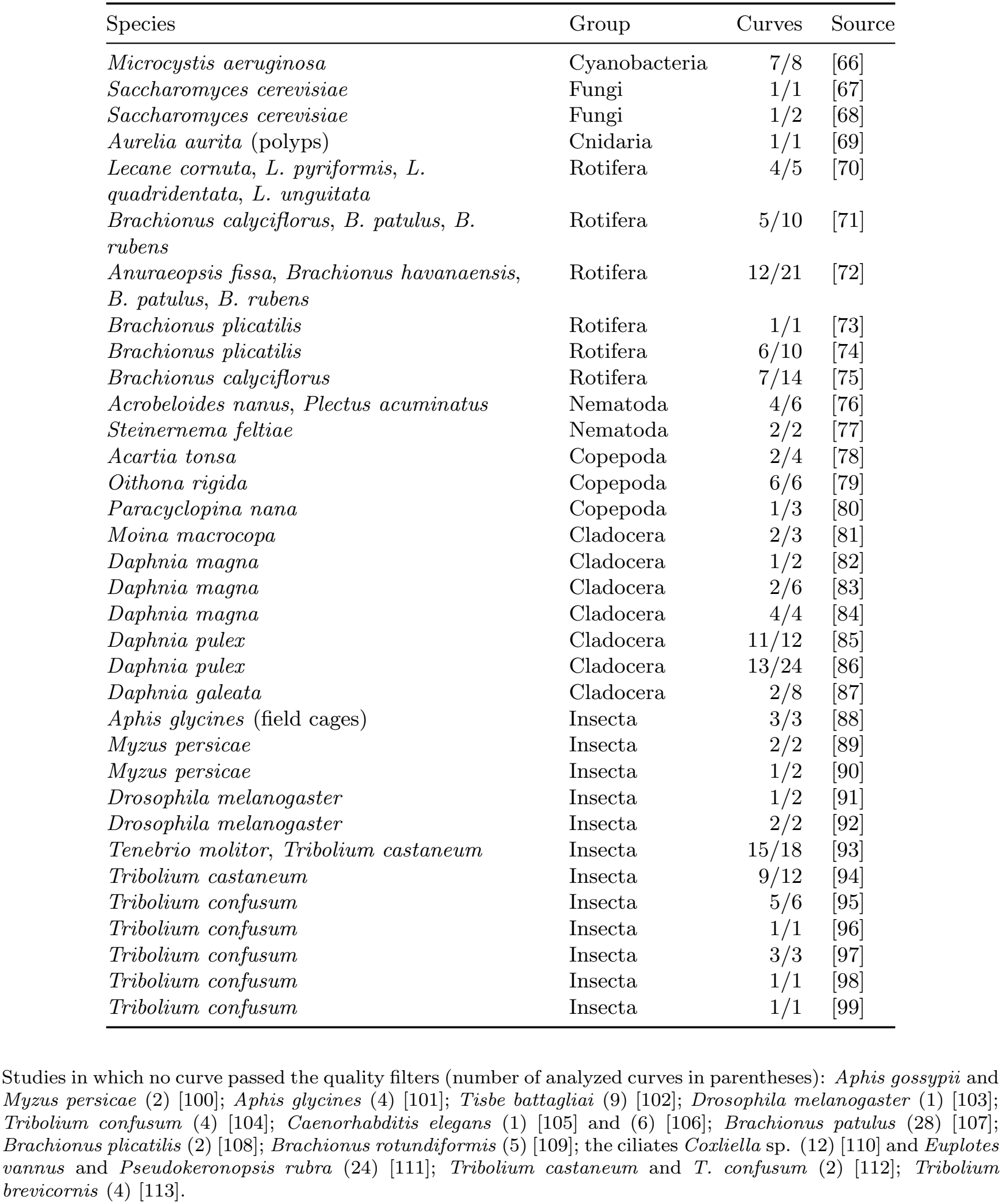
Studies composing dataset I. Only studies contributing at least one retained curve are listed, with the species of the retained curves. Curves are given as retained/analyzed.

| Species | Group | Curves | Source |
|---|---|---|---|
| *Microcystis aeruginosa* | Cyanobacteria | 7/8 | [66] |
| *Saccharomyces cerevisiae* | Fungi | 1/1 | [67] |
| *Saccharomyces cerevisiae* | Fungi | 1/2 | [68] |
| *Aurelia aurita* (polyps) | Cnidaria | 1/1 | [69] |
| *Lecane cornuta*, *L. pyriformis*, *L. quadridentata*, *L. unguitata* | Rotifera | 4/5 | [70] |
| *Brachionus calyciflorus*, *B. patulus*, *B. rubens* | Rotifera | 5/10 | [71] |
| *Anuraeopsis fissa*, *Brachionus havanaensis*, *B. patulus*, *B. rubens* | Rotifera | 12/21 | [72] |
| *Brachionus plicatilis* | Rotifera | 1/1 | [73] |
| *Brachionus plicatilis* | Rotifera | 6/10 | [74] |
| *Brachionus calyciflorus* | Rotifera | 7/14 | [75] |
| *Acrobeloides nanus*, *Plectus acuminatus* | Nematoda | 4/6 | [76] |
| *Steinernema feltiae* | Nematoda | 2/2 | [77] |
| *Acartia tonsa* | Copepoda | 2/4 | [78] |
| *Oithona rigida* | Copepoda | 6/6 | [79] |
| *Paracyclopina nana* | Copepoda | 1/3 | [80] |
| *Moina macrocopa* | Cladocera | 2/3 | [81] |
| *Daphnia magna* | Cladocera | 1/2 | [82] |
| *Daphnia magna* | Cladocera | 2/6 | [83] |
| *Daphnia magna* | Cladocera | 4/4 | [84] |
| *Daphnia pulex* | Cladocera | 11/12 | [85] |
| *Daphnia pulex* | Cladocera | 13/24 | [86] |
| *Daphnia galeata* | Cladocera | 2/8 | [87] |
| *Aphis glycines* (field cages) | Insecta | 3/3 | [88] |
| *Myzus persicae* | Insecta | 2/2 | [89] |
| *Myzus persicae* | Insecta | 1/2 | [90] |
| *Drosophila melanogaster* | Insecta | 1/2 | [91] |
| *Drosophila melanogaster* | Insecta | 2/2 | [92] |
| *Tenebrio molitor*, *Tribolium castaneum* | Insecta | 15/18 | [93] |
| *Tribolium castaneum* | Insecta | 9/12 | [94] |
| *Tribolium confusum* | Insecta | 5/6 | [95] |
| *Tribolium confusum* | Insecta | 1/1 | [96] |
| *Tribolium confusum* | Insecta | 3/3 | [97] |
| *Tribolium confusum* | Insecta | 1/1 | [98] |
| *Tribolium confusum* | Insecta | 1/1 | [99] |

Studies in which no curve passed the quality filters (number of analyzed curves in parentheses): *Aphis gossypii* and *Myzus persicae* (2) [100]; *Aphis glycines* (4) [101]; *Tisbe battagliai* (9) [102]; *Drosophila melanogaster* (1) [103]; *Tribolium confusum* (4) [104]; *Caenorhabditis elegans* (1) [105] and (6) [106]; *Brachionus patulus* (28) [107]; *Brachionus plicatilis* (2) [108]; *Brachionus rotundiformis* (5) [109]; the ciliates *Coxliella* sp. (12) [110] and *Euplotes vannus* and *Pseudokeronopsis rubra* (24) [111]; *Tribolium castaneum* and *T. confusum* (2) [112]; *Tribolium brevicornis* (4) [113].

## Notes

https://doi.org/10.5281/zenodo.20200296

### Competing Interest Statement

The authors have declared no competing interest.

### Summary of Updates

Revised title and abstract. Materials and Methods moved back to the main text. Small changes in the main text. Updated θ inference (accounting for discrete sampling) on a curated dataset of 3,995 curves; Fig. 2 updated accordingly. New: Fig. S6 (per-curve profile-likelihood confidence intervals for θ), Fig. S9 (parameter-free prediction of the E. coli growth-rate profile at 7.5 mM glycerol), Table S1 (datasets B to I) and Table S2 (studies composing dataset I). Former Figs. S6 and S7 are now Figs. S7 and S8.

